# Assessment and Optimization of Collective Variables for Protein Conformational Landscape: GB1 *β*-hairpin as a Case Study

**DOI:** 10.1101/333047

**Authors:** Navjeet Ahalawat, Jagannath Mondal

**Affiliations:** Tata Institute of Fundamental Research, Center for Interdisciplinary sciences, Hyderabad 500107, India

**Author notes:** Electronic mail (Jagannath Mondal).

## Abstract

Collective variables (CV), when chosen judiciously, can play an important role in recognizing rate-limiting processes and rare events in any biomolecular systems. However, high dimensionality and inherent complexities associated with such biochemical systems render the identification of an optimal CV a challenging task, which in turn precludes the elucidation of underlying conformational landscape in sufficient details. In this context, a relevant model system is presented by 16residue, *β* hairpin of GB1 protein. Despite being the target of numerous theoretical and computational studies for understanding the protein folding, the set of CVs optimally characterizing the conformational landscape of, *β* hairpin of GB1 protein has remained elusive, resulting in a lack of consensus on its folding mechanism. Here we address this by proposing a pair of optimal CVs which can resolve the underlying free energy landscape of GB1 hairpin quite efficiently. Expressed as a linear combination of a number of traditional CVs, the optimal CV for this system is derived by employing recently introduced Time-structured Independent Component Analysis (TICA) approach on a large number of independent unbiased simulations. By projecting the replica-exchange simulated trajectories along these pair of optimized CVs, the resulting free energy landscape of this system are able to resolve four distinct well-separated metastable states encompassing the extensive ensembles of folded,unfolded and molten globule states. Importantly, the optimized CVs were found to be capable of automatically recovering a novel partial helical state of this protein, without needing to explicitly invoke helicity as a constituent CV. Furthermore, a quantitative sensitivity analysis of each constituent in the optimized CV provided key insights on the relative contributions of the constituent CVs in the overall free energy landscapes. Finally, the kinetic pathways connecting these metastable states, constructed using a Markov State Model, provide an optimum description of underlying folding mechanism of the peptide. Taken together, this work offers a quantitatively robust approach towards comprehensive mapping of the underlying folding landscape of a quintessential model system along its optimized collective variables.

## I. INTRODUCTION

The complexity of protein folding requires one to employ a multi-dimensional approach to understand the diversity of folding pathways via transition states ^1^. Over the last decade, computer simulation, with the advent of new algorithms and better hardware, has slowly gained its reputation as a suitable method for exploring protein folding. While Molecular Dynamics (MD) simulations, by harnessing the power of special purpose computers, have recently lived up to its true potential of exploring the complete protein folding process in an unbiased fashion ^2^, enhanced sampling schemes have also emerged as efficient and costeective methods for simulating the protein folding landscape. The proven ability of these enhanced sampling schemes to overcome multiple, deep rugged free energy minima within modest simulation time have made them popular techniques among the scientific communities. However, the key to the quantitative analysis of protein folding lies in exploring the underlying conformational transition along one or multiple collective variables (CV). More over, majority of these enhanced sampling schemes (such as metadynamics ^3–5^ and umbrella sampling^6^) require choosing an appropriate collective variable (CV) which describes the progress of the conformational transition. Hence, the quality, reliability, and usefulness of the explored conformational ensemble depend on the quality of the chosen CV. As key requirement, a chosen CV should capture all the relevant slowly changing degrees of freedom in the system and should separate out all relevant metastable states. However, the lack of an unique approach for dimensionality reduction has made the quest for an appropriate collective variable for any physical process, including protein folding, an emerging problem.

Identification of optimal CV in very complex biological system is currently being recognized as a true chalenge which has motivated several methodological developments. Tiwary and Berne have recently proposed ^8^ spectral gap optimization of order parameters (SGOOP) algorithm which identifies the slowest linear or nonlinear combination of CVs using maximum caliber-based approach. Sultan and Pandey^9^ have also introduced another CV optimization method coined as “TICA-MetaD” which identifies the linear combinations of CVs using the time-structure based independent component analysis (TICA) approach^10–14^. Very recently, Parrinello and coworkers ^15,16^ also have developed an alternate approach to both SGOOP and TICAMetaD, namely VACMetaD. In VAC-MetaD, a trial combination of conventional CVs is first employed to carry out biased simulations, from which few slow modes (based on spectral gap in timescales) are variationally identified using TICA approach. Finally, these slow modes are used as new CVs to perform further well tempered metadynamics. All these methods, while retaining mutual distinction, have shown that optimized CVs, derived using these methods, can guide the sampling in a remarkable way. Taking cue from this, in this work, by using a TICA based approach, we put forward the proposal of an optimized set of CV for exploring conformational landscape of GB1 hairpin, a classic model system for understanding protein folding.

**FIG. 1:**
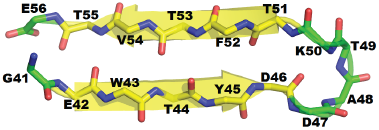
Representation of the crystal three-dimesnsional stucture of GB1 hairpinpeptide (PDB ID 1GB1^7^). Only backbone atoms for every residues are shown. The cartoon representation is also overlaid using yellow color for beta-sheet and green color for loop regions.

The system of our interest is the C-terminal domain of GB1 protein^7^ which forms a, *β*-hairpin (Figure 1) and has been extensibly characterized by both experiments and simulations. It has been observed that it independently folds into, *β*-hairpin structure which is similar to its native state. Although it is a fast folder (with folding time ∼6 *μ*s) and has 50% folded fraction at 300 K, there is no consensus on the underlying folding mechanism of this peptide. The folding mechanism of this peptide has been previously investigated using Replica-Exchange Molecular Dynamics (REMD) simulations^17^ in explicit solvent that performs a random walk in temperature space and allows system to overcome the energy barriers.^18–21^ By projecting simulated data along the conventional and intuitively chosen CVs, such as Radius of gyration (Rg), rootmeansquared deviation (RMSD), fraction of native contacts (Qn), and native state hydrogen bonds (Qh), these works reported the presence of three metastable states in GB1. The first two dimensions of principal component analysis (PCA) of REMD data also described only three free energy basins^22^. Previous metadynamics simulations studies of, *β-*hairpin peptide^23–25^ used the conventional collective variables such as Rg and native hydrogen bonds to describe and accelerate the folding process. These studies also reported that this peptide has Lshaped free energy landscape with maximum three metastable states. On the other hand, structure based clustering^19^ and the sketch-map analysis^26^ of simulated data suggest that this peptide could have more than four different types of metastable conformations i.e, unfolded, collapsed, helical, and, *β*sheet structures. But none of the CVs, on its own, was able to distinguish all of these conformationally distinct states with proper energy-barrier. In these contexts, our current work provides a quantitative solution towards deriving an optimized CV for GB1 hairpin, which is capable of resolving the free energy landscape into well-separated basins.

In this work, we have employed a quantitative approach, based upon TICA^10–14^, to construct a set of optimized CVs of GB1 hairpin and in the process have elucidated the complete free energy landscape of this peptide via projection of REMD trajectories along these CVs. As will be presented in the current article, freeenergy landscape plotted against first two slowest varying TICs, obtained by projecting them along a linear combination of five traditional CVs, are able to resolve four distinct metastable states which are not otherwise visible using conventional CVs. Apart from recovering the folded,unfolded and molten globule ensembles of GB1, *β*hairpin, the free energy projection along optimized CVs were able to spontaneously recover a partial helical state of this protein, without needing to explicitly invoke helicity as a constituent CV, unlike previous works. Furthermore, the relative contributions of each constituent in the optimized CV were also quantitatively assessed. Finally, the kinetic pathways connecting the metastable ensembles were obtained under the framework of a Markov state Model, which provided a streamlined view of the underlying folding mechanism of GB1 hairpin.

## II. MATERIAL AND METHODOLOGY

### A. System preparation and molecular dynamics

The Cterminal domain of GB1 hairpin (residues 41–56) was taken out from the full-length GB1 protein NMR structure (PDB: 1GB1^7^). The peptide was capped with acetyl group on N-terminal and methyl amide on Cterminal. The 16residue long peptide Ace41GEWTYDDATKTFTVTE^56^Nme was modeled using AMBER03STAR^27,28^ forcefield and solvated in a truncated octahedron simulation cell containing 984 TIP3P^29^ water molecules. The system was charge-neutralized by using six Na^+^ ions, and three Cl^-^ ions. All simulations were performed using GROMACS 5.1^30^ package. The system was minimized for 10000 steps using steepest decent algorithm. Particle Mesh Ewald summation method^31^ was used for longrange electrostatic interactions with a 0.12 nm grid spacing and 1.0 nm cutoAll the bonds connected with H atoms were constrained using LINCS^32^ method. SETTLE algorithm^33^ was used to constrain the bonds and the angle of water molecules. During equilibration period, solvent molecules were first allowed to relax by restraining the positions of all heavy atoms of the peptide in NVT ensemble at 300 K using Berendsen^34^ thermostat for 200 ps. Then, the full system was equilibrated at same temperature without any restrain using velocity-rescaling^35^ method with a coupling constant of 0.5 ps for 500 ps. The system was further equilibrated in NPT ensemble at 300 K and 1 bar pressure for 5 ns using first Berendsen^34^ barostat, followed by Parrinello-Rahman^36^ barostat with a coupling constant of 2 ps. The equilibrated structure obtained at the end of the NPT simulation was used for all subsequent NVT simulations.

**FIG. 2:**
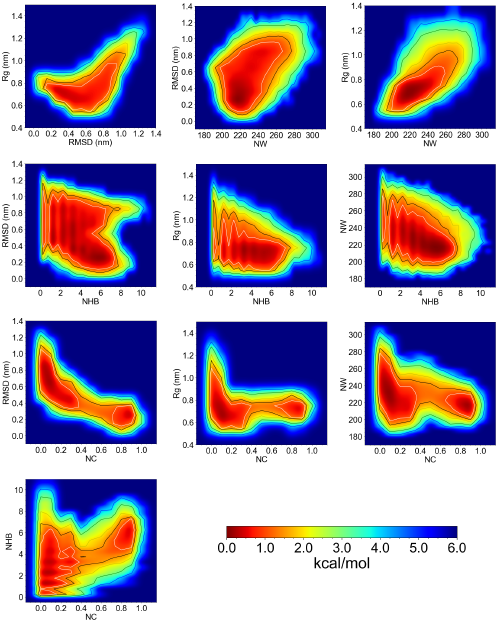
Two dimensional free energy surface along the conventional CVs: Rg, RMSD, Fraction of native contacts(NC), number of water around peptide(NW), and number backbone-backbone hydrogen bonds (NHB). Contours are drawn at 1(white),1.5(black), 2(yellow), 3(green), 4(blue) kcal/mol.

In this work, two types of simulations have been carried out: REMD simulations^17^ and multiple unbiased roomtemperature simulations. REMD simulation is first utilized to efficiently sample the conformational landscape and generate key conformations (by clustering) for initializing subsequent multiple unbiased room temperature simulations. Subsequently, 200 multiple unbiased simulations, spawned from REMD-derived snapshot, were performed and later used for deriving the optimized CVs using TICA model. The REMD-derived trajectories were finally employed again to project the conformational free energy landscape along the optimized CV.

For the purpose of REMD simulations, a total of 32 replicas were used with temperature range 278-595 K. The temperatures used were as 278, 287, 295, 303, 312, 321, 329, 338, 346, 355, 365, 375, 385, 396, 406, 416,427, 437, 448, 459, 470, 482, 493, 505, 517, 528, 539,551, 562, 573, 584, and 595 K. Each replica was initially equilibrated at the given temperature in the NVT ensemble for 20 ns. Subsequently, the simulations were carried out for 400 ns per replica with a time step 2 fs, and the exchange between adjacent replica were attempted at an interval of every 10 ps. The coordinates were saved every 2 ps for the purpose of further analysis. Parallel Tempering Weighted Histogram Analysis Method (PTWHAM)^37^ was employed for the calculation of free energies. Figure S1A and S1B in SI text shows the time evolution of two representative replicas of simulations. A good exchange among replicas during the course of the simulation demonstrated that each trajectory performs a random walk in temperature space. Furthermore, the distributions of the potential energy for all of the replicas (Figure S1C) clearly show reasonable overlap between adjacent replicas and guarantee reasonable exchange probabilities. The good correlation between calculated ensemble averaged NMR chemical shift using SPARTA+^38^ at 278 K and experimental chemical shift^39^ (Figure S2) further ensures that simulation is well-converged and equilibrium ensemble is properly sampled.

The REMD conformations at 303 K were subsequently divided into 200 clusters using the RMSD based kmeans clustering algorithm^40^. Then one conformation from each of the clusters was selected randomly and used as a starting conformation for subsequent unbiased MD simulations. Each trajectory was simulated up to 100 ns at 303K with the same setup as used for REMD simulations.

### B. Timestructured independent component analysis

The Independent Component Analysis (ICA)^41^ is a statistical and computational technique for transforming an observed multidimensional data into statistically independent components, called the independent components (ICs), and has been used extensively in the field of signal processing and data analysis. Although there are various ICA methods, we have employed the Time-structure based Independent Component Analysis (TICA) which is a kinetically motivated unsupervised learning method to find the slowest degrees of freedom without losing important kinetic information. In TICA, the goal is to find a linear combination of input coordinates or structural order parameters that maximizes the autocorrelation of transformed coordinates at a particular lag time. The complete mathematical derivation for the TICA formalism could be found in the references^10–14^. Here, we will briefly describe it. It should be noted that, for clarity, most of our notations is borrowed from the references^9,14^.

Let 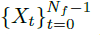 be a multidimensional time series MD data where each frame of trajectory is represented via a column vector of ddimensions for a system. The *X*_*t*_ is a frame at time t and *N*_*f*_ is the total number of frames in MD trajectory data. Further, we are assuming that the mean of data is zero. The goal of TICA is to maximize the function *f*(|*v*⟩),which is autocorrelation function of the projection of |*X*_*t*_⟩ onto |*v*⟩

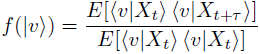

Here, τ some lag time greater than zero. Since the inner product is symmetric, we can rewrite this equation as:

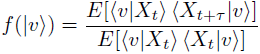

The outer products *E*[*X*_*t*_*X*_*t+τ*_] and *E*[*X*_*t*_*X*_*t*_] in the above equation are the same as the timelag correlation matrix and *C* ^(*τ*)^the covariance matrices Σ, respectively. Hence, we can rewrite the function as:

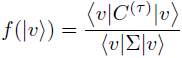

After applying the constraint on TICs to have unit variance: ⟨*v*|Σ|*v*⟩ =1 and setting up optimization problem that is similar to PCA and solvable via Lagrange multipliers methods, it can be shown that the solution to above equation is also a solution to the generalized eigenvalue problem:

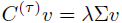

Further, the remaining TICs, the slowest projections, can be obtained similarly by constraining the TICs to be uncorrelated with the previously found solutions.

To find the desired TICs, we have followed the procedure described in references^9,14^ with the following steps: (1)We have run 200 unbiased (100 ns each) MD simulations starting from the conformations obtained after the clustering of 303 K REMD data, (2) then, five dimensional time series data of five traditional and frequently used CVs, namely, Rg of all backbone heavy atoms, RMSD of all backbone heavy atoms with respect to the crystal structure, fraction of native contacts (NC), the number of backbone-backbone hydrogen bonds(NHB), and the number of water molecules around 5 Angstrom of peptide (NW) were used as input to build the TICA model (which is a linear combination of the input CVs) with a lag time of 1 ns using MSMBuilder3 package^42^. The steps to build TICA models are:

- Compute time-lag correlation matrix, *C*^(*τ*)^, and covariance matrix, Σ, from multidimensional timeseries data.
- Symmetrize the *C*^(*τ*)^.
- Solve the *C*^(*τ*)^*v* = λΣ*v*.
- Then select desired number of eigenvectors (TICs) with the top (largest) eigenvalues.

In this work, two TICs with the largest eigenvalues (named hereafter as“TIC1” and “TIC2”) were used in all the analysis.

Thus, the resulting TIC1 and TIC2 are linear combinations of five input CVs and can be represented as (see equation 1):

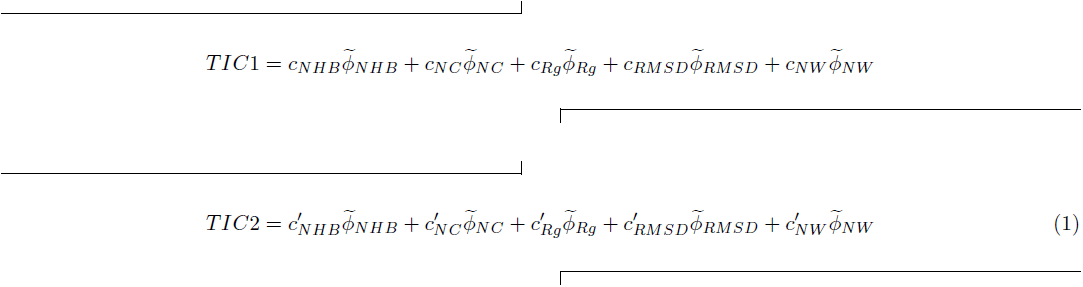

where 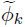 represents kth mean free input CV and *c*_*k*_ and 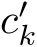 are associated coefficients for TIC1 and TIC2 respectively. All the input CVs were first rescaled between 0 and 1 before using them to build the TICA model.

Toward this end, NC and NHB were calculated using a switching function;

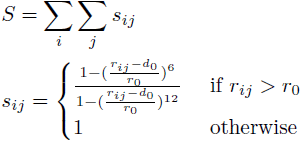

where *d*_0_ = 0.05 nm, and *r*_sub>0_ = 0.65 nm were used for NC calculation and *d*_0_ = 0.1 nm, and *r*_0_ = 0.23 nm were used for the NHB calculation with D MAX = 0.25 nm as implemented in plumed^43^. *r*_*ij*_ is a distance where i and j are pairs of all heavy atoms in native state within 0.65nm whereas in case of NHB i and j are pairs of O and HN atoms of backbone.

## III. RESULTS AND DISCUSSIONS

### A. FreeEnergy Landscape of the, *β*hairpin along traditional CVs does not resolve all important states

GB1 hairpin has remained a classic model system for investigating the protein folding. Majority of the previous works have explored the conformations of, *β*hairpin equilibrium by projecting the structural ensemble along several conventional CVs, intuitively chosen based on inspection of simulation trajectories. The extensive studies on this system have zeroed in on five CVs, namely Rg, RMSD, fraction of native contacts (NC), the number of backbone-backbone hydrogen bonds(NHB), and the number of water molecules around 5 Angstrom of peptide (NW). Before we arrive at our optimized sets of CVs for GB1 hairpin, we first assessed the underlying free energy landscape by projecting our extensively performed (total 32 replicas from 278 to 595 K, each of 400 ns) REMD simulated conformational ensembles along these popular and conventional CVs, taking a pair of conventional CVs at a time. Figure 2 shows the twodimensional free-energy surfaces as a function of pair of CVs, at 303 K. These multi-dimensional free energy surfaces projected along the conventional CVs are very similar to the previous works and suggest that there exist only two metastable states in the conformational landscape of GB1 hairpin. Most of the earlier studies, focusing on the folding mechanism of the beta-hairpin have attempted to derive a CV that can distinguish between two major events i.e., collapsing event and hydrogen bond formations. The culmination of all previous works have majorly pointed towards Radius of gyration (Rg) and the number of intramolecular backbone-backbone hydrogen bonds (NHB) of the GB1 hairpin as potentially capable of serving as better CVs. Accordingly, most of the folding mechanisms of GB1 hairpin have revolved around the projection of free energy profiles along these two CVs. Previous studies have shown that the free energy landscape as a function of Rg and native hydrogen bonds is Lshaped with either two states^24^ (folded and unfolded) or three states^22,25,26^ (folded, collapsed, and unfolded). When projected along same CVs, our results find it to be a two state folder (Figure 2). However, none of the them is able to show the importance of helix formation during folding process of a beta-hairpin peptide. To understand the role of helical structures in the folding process, previous investigations explicitly needed^44,45^ to use the fraction of helical contacts as one of the CVs to separate out the helical structures from unfolded ensemble. The overall analysis indicates the lack of an well-constructed and rationally optimized CV which holds the potential of automatically resolving the free energy landscape of GB1 hairpin into all possible basins and can demarcate the helical structures from unfolded ensemble without taking recourse to explicit usage of any secondary structure analysis.

### B. Optimized CV for GB1 based on TICA resolves free energy landscape into four metastable states

The aforementioned discussion demonstrated that these conventional CVs, when used as a pair, are not sufficient to resolve all the relevant metastable states of GB1 hairpin. In these contexts, as detailed in the method section, we propose a pair of optimized CVs, namely TIC1 and TIC2 (see equation 1), derived using a TICA-based approach by sampling over 200 unbiased trajectories, each 100 ns long. As shown in equation 1, the optimized CVs are the linear combination of five conventional CVs, namely Rg, RMSD, NC, NHB, and NW. Projection of unbiased trajectories along slowest first dimension TIC1 is shown in Figure S3 which suggests that the peptide has explored enough phase space with sufficient exchange between different conformations along this optimized CV. Furthermore, Table S1 presents the coefficients associated with the each of five constituent conventional CVs. We find that the values of these coefficients are robust against variation of trajectory length for construction of TICA dimensions.

The REMD trajectory at 303 K was projected along the optimized CVs TIC1 and TIC2 (Figure 3A and B). Figure 3B shows the two dimensional free energy surface along TIC1 and TIC2. We find that the system shows it’s characteristic Lshaped free energy landscape with four free-energy basin named as ms1, ms2, ms3 and ms4 (Figure 3B). After clustering the conformations belonging to those minima, we identify that conformations of minima ms1 and ms4 belong to the unfolded and the folded state whereas ms2 and ms3 are collapsed molten globule intermediate states. Interestingly, the state ms2 represents the conformations with helical contents. These molten globule states with partial helicity were not easy to distinguish using traditional CVs. The previous study needed to use helicity as one of the reaction coordinates to identify them in free energy landscape. However, the CVs, optimized in the current work, can resolve the basin with partial helical structure automatically. The one dimensional free energy surface along slowest TIC1 shows two dominated unfolded (ms1) and folded (ms4) states with very shallow intermediate states ms2 and ms3 (Figure 3A). The two intermediate metastable states ms2 and ms3 are not prominent enough along first slowest direction but can be easily distinguished after considering first two TIC dimensions. Thus, we suggest using more than one dimension to get free energy surface with proper metastable states in case of GB1 hairpin.

**FIG. 3:**
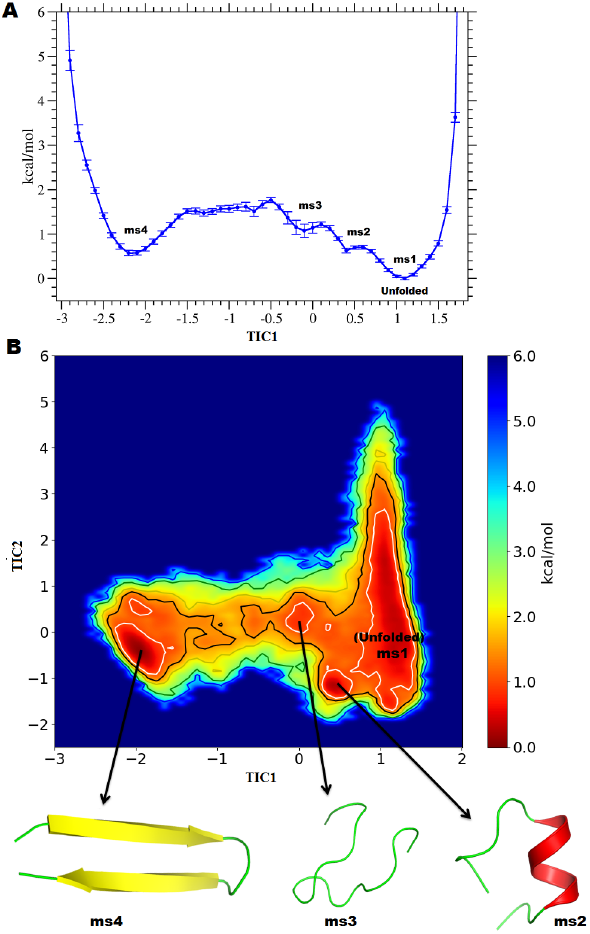
(A) One dimensional free energy surface along the first slowest TIC1 dimension, (B) Two dimensional free energy surface along the first two slowest TICA dimensions: TIC1 and TIC2. Representative structures belonging to a particular minima are also shown. Contours are drawn at 1(white),1.5(black), 2(yellow), 3(green), 4(blue) kcal/mol.

### C. Assessment of key contributors in the optimized CV

One of the major benefits of using feature space instead of coordinate space in the optimized CV is that these features are easy to implement in enhanced sampling method and more importantly one can quantitatively assess the role of each function in the underlying mechanism. As illustrated by the coefficients of optimized CV (Figure 4) (normalized for comparison), in our case, the native contacts of peptide (NC) contribute the most towards the construction of slowest dimension TIC1first slowest TIC1 dimension, (B) Two dimensional free TIC1 and TIC2. Representative structures belonging to 1(white),1.5(black), 2(yellow), 3(green), 4(blue)whereas Rg contributes more towards second slowest dimension TIC2. To assess the effect of each constituent CV in free energy landscape, we perform a negative control analysis, where we systematically remove one of the constituent CVs from the optimized CVs, one at a time. Subsequently, same procedure, as previously described, is iterated to obtain the optimized CV using remaining four conventional CVs. Figure 5 shows the effect of absence of each constituent CV on the two-dimensional free energy landscape. As is evident from the figure 5, optimization of CV in absence of NC and Rg triggers largest change in the shape of the free energy landscape, whereas absence of other constituent CVs has minimal affect on the overall free energy landscape. Thus, both weights of constituent CVs in the optimized CV and the cross validation via systematically removing each of them, one at a time, provide key insights on their effective contributions on the optimized CVs.

**FIG. 4:**
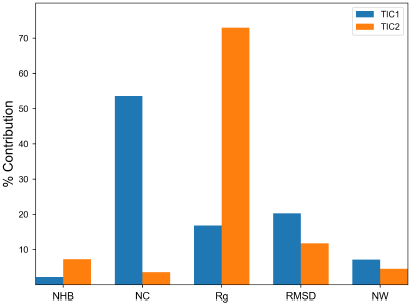
Percent contributions of the conventional input CVs to the first two slowest dimensions (TIC1and TIC2) are shown as a bar plot. Here, only absolute value of coefficients (see Table S1 in SI text) is considered for contribution calculations.

Figure 4 indicates that even though NC and Rg contribute the most towards TIC1 and TIC2 respectively, the other input CVs also have considerable contributions in clearly resolving the four metastable states shown in Figure 3. This is physically corroborated by figure S4 (see SI) which illustrates the values of each input CVs averaged along twodimensional grids of TIC1 and TIC2. As we see, while values of NC change sharply along TIC1, suggesting the major change of nativeness occurring along TIC1, it alone can not differentiate between partially folded helical (ms2) and collapsed (ms3) structures shown in Figure 3. Similarly, while Rg values change sharply along TIC2, it also can not differentiate between fully folded (ms4) and partially helical structure(ms2). These emphasize the significantt role of the other input CVs as a linear combination in resolving the whole free energy landscape of GB1 hairpin.

### D. Markov state Model along optimized dimensions kinetically connects GB1 folding pathways

GB1 hairpin has been the subject of extensive Markov state model (MSM) based investigations for obtaining the kinetics of folding and unfolding of this peptide.^45–50^To check or validate the robustness of the our TICA based optimized CVs, we have generated the MSMs from unbiased short trajectories by considering only first twoTICs at dierent lag time and compared the relaxation timescale of slowest process with previous works. Figure 6A shows the relaxation time (implied timescales)of ten slowest processes of this peptide. Our CVs also show same relaxation timescale range and similar associated gap between them as in previous studies^46,50^. The first large gap in the spectrum of the relaxation times is between first and second mode (top blue line) whereas the second gap is then between third and fourth slow modes (green line from top), which suggests that there are maximum four metastable states in the underlying free energy landscape. Further, we have chosen 5 ns as lag time to build the final MSM and coarse grained it into four metastable states using Robust Perron Cluster Analysis (PCCA+)^51,52^. Interestingly, these four metastable states obtained from hidden MSM correspond to the energy minima basin (ms1-4) obtained using REMD data. Furthermore, we used transition path theory (TPT)^53–55^ as implemented in PyEMMA^56^ package to identify the folding paths from unfolded state. The TPT analysis suggests that transition from unfolded state (ms1) to folded state (ms4) is more preferable through intermediate ms3 than ms2 (helical state). The TPT analysis also demonstrates that helical intermediate state (ms2) first converts into ms3 to achieve final native state (ms4) (see figure 6 B). Previous kinetic studies also have suggested that the folding rate of this peptide is very slow if the starting state is helical. Thus, the combinations of different input CVs improve the geometrical classification of different kind of conformations and TICA, which is a variational approach, helps in the identification of few slow linear combination of the input CVs to obtain proper metastable states with correct kinetics. Overall, these brief kinetic analysis based on only first two slowest CVs namely TIC1 and TIC2 highlight the usefulness of these reduced but optimum dimension to provide insights in the underlying folding mechanism of this peptide.

## IV. CONCLUSIONS

In summary, the current article demonstrates that the CVs optimized using linear combination of feature space with recently described TICA approach offers a promising direction towards exploring the free energy landscape of GB1 hairpin with properly resolved intermediates. By using feature space as the input CVs, the optimized CVs can uncover atomistic details of the free energy landscape of this quintessential model system for protein folding quite efficiently. Along with resolving the free energy landscape into four well-resolved metastable states of the short-chain peptide, the optimized CV could automatically recover a partial helical state of this system, without requiring the utilization of helicity as a constituent CV. As an added advantage of expressing the optimal CV in the form of linear combination of multiple conventional CVs, the method allows one to test the contribution of each constituent CVs in the overall free energy landscape of GB1 hairpin. These feature spaces with TICA approach are easy to implement in enhanced sampling method. More over, the MSMbased kinetic analysis based upon only a limited number of optimal CVs is found to yield crucial insights on the folding pathways of GB1 hairpin. Overall, we believe that this work provides a general and efficient approach to explore the protein folding problem and further it can be extended to understand the other complex biomolecular mechanisms and pathways.

**FIG. 5:**
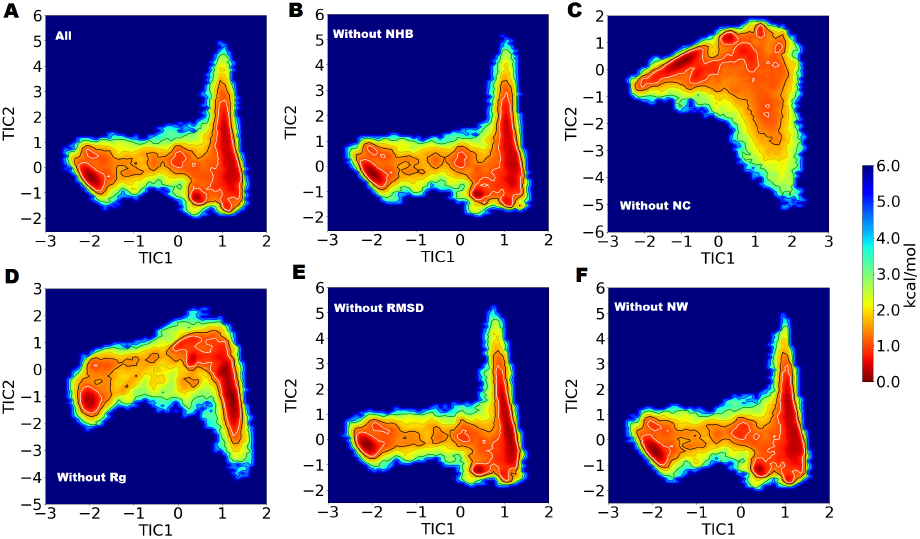
Two dimensional free energy surface along the first two slowest dimensions, (A) considering all the functions: NHB (number of hydrogen bonds), NC (fraction of native contacts), Rg (radius of gyration), RMSD (root-mean-square deviation), NW (number of water molecules around peptide), after dropping (B) NHB, (C) NC,(D) Rg, (E) RMSD, (F) NW from the optimal CV. NC and Rg show largest effect on free energy landscape whereas others show minimal effect. Contours are drawn at 1(white),1.5(black), 2(yellow), 3(green), 4(blue) kcal/mol.

**FIG. 6:**
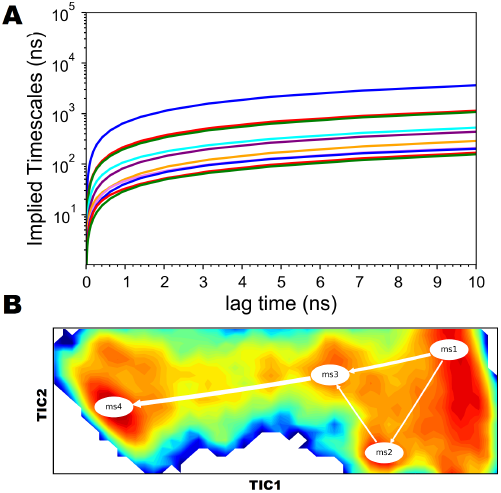
Top ten implied timescale values as a function of lag time. Different color line represents the different implied timescales (relaxation time). Transition pathways between unfolded metastable state to folded state (obtained using Transition Path Theory) shown on the top of free energy landscape obtained from REMD data. elliptical shape represents the metastable state whereas thickness of arrow represents the net flux going from one state to another state.

Finally, we note that while our results illustrate the merits of the using the combination of physically meaningful CVs with TICA approach, the computational cost and quality of the optimized CVs can still be improved by incorporating the nonlinearity in the combination. In the last few years, the deep learning, a particular variant of machine learning approach, is emerging as an efficient approach for identification of linear and/or nonlinear reaction coordinates to perform biased sampling^57–60^. In future, it would be interesting to compare the behavior (i.e. weight) of these input feature variables in machine learning approach to decode the inherent complexity of protein folding processes.

## ACKNOWLEDGMENTS

This work was supported by computing resources obtained from shared facility of TIFR Hyderabad, India. JM would like to acknowledge intramural research grants obtained from TIFR, India and Ramanujan Fellowship of DST, India. Part of the work was carried out in San Diego supercomputing resources provided by XSEDE (TGCHE150024).

## V. SUPPLEMENTAL INFORMATION

Figures of Time profiles of multiple replica, probability distributions of energetics of replica, comparison of simulated and experimental chemical shifts of GB1 hairpin, time profiles of slowest optimal CV TIC1, dependences of all CVs on TIC1 and TIC2 coordinates, Table containing coefficients of different constituent CVs. (PDF).

## Supporting Information for “Assessment and Optimization of Collective Variables for Protein Conformational Landscape: GB1 β-hairpin as a Case Study”

Navjeet Ahalawat and Jagannath Mondal*

Tata Institute of Fundamental Research, Center for Interdisciplinary Sciences, Hyderabad, India 500107

**Figure S1:**
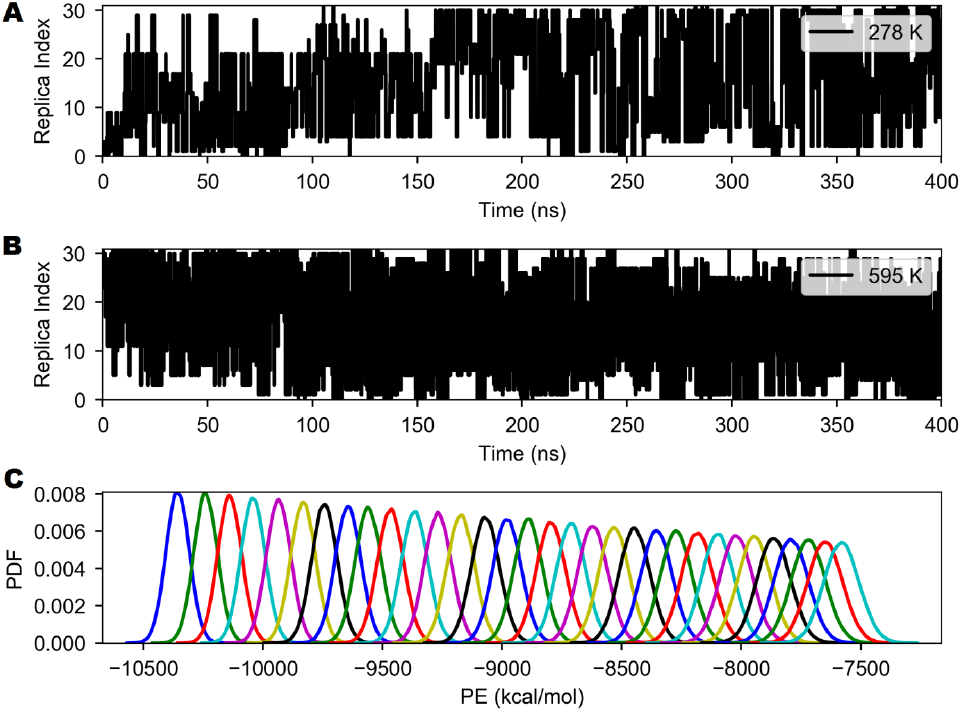
Time series of exchange among replicas at three representative temperatures (A) 278 K, (B) 595 K. (C) The canonical probability distributions of the total potential energy for 32 temperatures.

**Figure S2:**
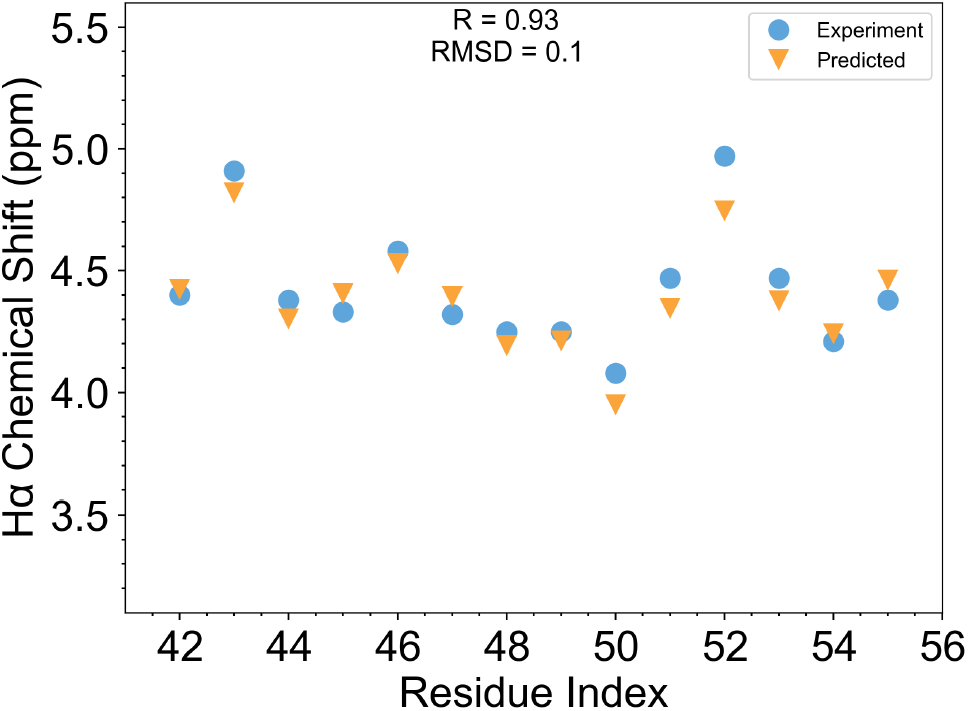
Average chemical shifts of Hα atoms of each residue are shown which were calculated from REMD simulated ensemble (considering all the frames) at 278 K using the program SPARTA+. Our results show a very good correlation between the calculated (predicted) and experimental chemical shift with R = 0.93 and root-mean-square deviation RMSD = 0.1 ppm with respect to experimental value

**Figure S3:**
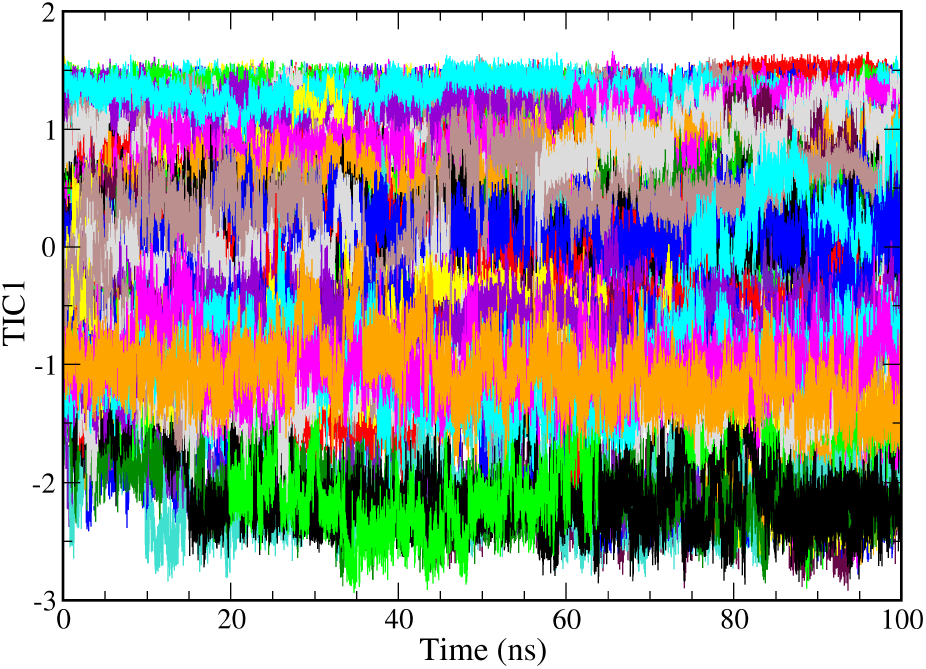
Time series plot of a few out of 200 short trajectories along the slowest dimension TIC1 to demonstrate that there is enough exchange between different conformations in the phase space.

**Figure S4:**
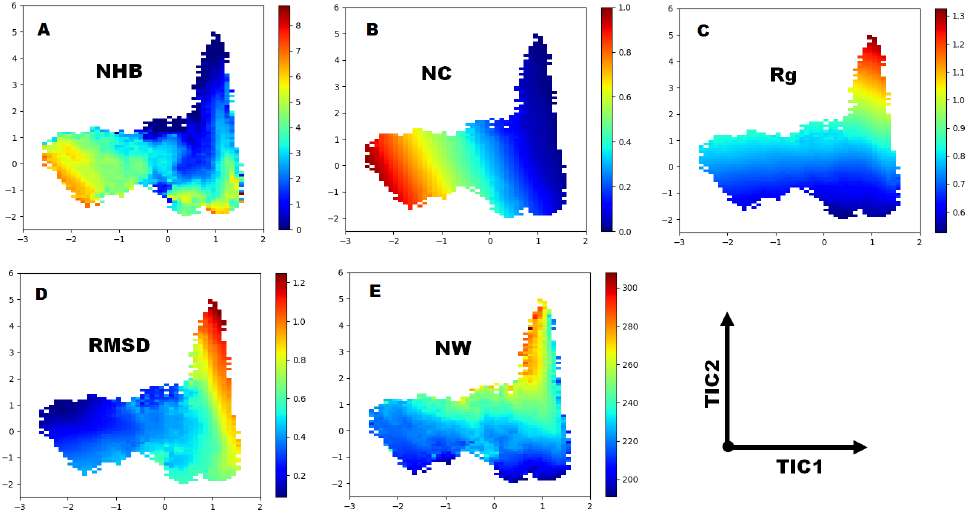
The dependence of all the CVs on TIC1 and TIC2 coordinates. Here, each conformation of REMD simulation at 303 k is represented by a point in the TIC1-TIC2 plane. Different conformations are represented by the color determined by the value along that particular input CVs (NHB, NC, Rg, RMSD, and NW). (A) Distribution of all the conformations are shown in the form of number of hydrogen bonds along TIC1 and TIC2 where blue color represents less number of hydrogen bonds and red mean high number of hydrogen bonds.Similarly, (B) fraction of native contacts, (C) Radius of gyration (Rg) which range between 0.5 nm to 1.3 nm, (D) RMSD (nm), and (E) number water molecules within 5 Å of peptide

**Table S1:** TIC coefficients calculated considering different lengths of short trajectories.

| TIC1 Coefficients |  |  |  |  |  |
| --- | --- | --- | --- | --- | --- |
| Time (ns) | $C_{\text{NHB}}$ | $C_{\text{NC}}$ | $C_{\text{Rg}}$ | $C_{\text{RMSD}}$ | $C_{\text{NW}}$ |
| 100 | 0.1403 | -3.4493 | -1.0840 | 1.3081 | -0.4593 |
| 90 | 0.1461 | -3.4567 | -1.1044 | 1.3367 | -0.4566 |
| 80 | 0.1499 | -3.4559 | -1.1206 | 1.3718 | -0.4569 |
| 70 | 0.1585 | -3.4603 | -1.1240 | 1.3983 | -0.4589 |
| TIC2 Coefficients |  |  |  |  |  |
| Time (ns) | $C'_{\text{NHB}}$ | $C'_{\text{NC}}$ | $C'_{\text{Rg}}$ | $C'_{\text{RMSD}}$ | $C'_{\text{NW}}$ |
| 100 | -0.8187 | -0.4036 | 8.2601 | -1.3305 | 0.5111 |
| 90 | -0.8053 | -0.3980 | 8.2666 | -1.2606 | 0.5103 |
| 80 | -0.7785 | -0.3439 | 8.2907 | -1.0868 | 0.5028 |
| 70 | -0.7692 | -0.4030 | 8.3714 | -1.1432 | 0.4819 |

## References

1. C. R. Matthews, Annu. Rev. Biochem. 62, 653 (1993).

2. R. O. Dror, R. M. Dirks, J. P. Grossman, H. Xu, and D. E. Shaw, Annu Rev Biophys 41, 429 (2012).

3. A. Laio and M. Parrinello, Proc. Natl. Acad. Sci. U.S.A. 99, 12562 (2002).

4. A. Barducci, G. Bussi, and M. Parrinello, Phys. Rev. Lett. 100, 020603 (2008).

5. O. Valsson, P. Tiwary, and M. Parrinello, Annu Rev Phys Chem 67, 159 (2016).

6. G. Torrie and J. Valleau, Journal of Computational Physics 23, 187 (1977).

7. A. Gronenborn, D. Filpula, N. Essig, A. Achari, M. Whitlow, P. Wingfield, and G. Clore, Science 253, 657 (1991), http://science.sciencemag.org/content/253/5020/657.full.pdf.

8. P. Tiwary and B. J. Berne, Proc. Natl. Acad. Sci. USA 113, 2839 (2016).

9. M. M. Sultan and V. S. Pande, Journal of Chemical Theory and Computation 13, 2440 (2017).

10. G. P. Hernandez, F. Paul, T. Giorgino, G. D. Fabritiis, and Noe, The Journal of Chemical Physics 139, 015102 (2013).

11. F. Nuske, B. G. Keller, G. Perez Hernandez, A. S. J. S. Mey, and F. Noe, Journal of Chemical Theory and Computation 10, 1739 (2014).

12. R. T. McGibbon and V. S. Pande, The Journal of Chemical Physics 142, 124105 (2015).

13. R. T. McGibbon, B. E. Husic, and V. S. Pande, The Journal of Chemical Physics 146, 044109 (2017).

14. C. R. Schwantes and V. S. Pande, Journal of Chemical Theory and Computation 9, 2000 (2013).

15. J. McCarty and M. Parrinello, The Journal of Chemical Physics 147, 204109 (2017).

16. Y. I. Yang and M. Parrinello, Journal of Chemical Theory and Computation, doi:10.1021/acs.jctc.8b00231 PMID: 29715017, https://doi.org/10.1021/acs.jctc.8b00231.

17. Y. Sugita and Y. Okamoto, Chemical Physics Letters 314, 141 (1999).

18. J. Juraszek and P. G. Bolhuis, Journal of Physical Chemistry B 113, 16184 (2009).

19. R. B. Best and J. Mittal, Proteins: Structure, Function and Bioinformatics 79, 1318 (2011).

20. M. Andrec, A. K. Felts, E. Gallicchio, and R. M. Levy, Proceedings of the National Academy of Sciences of the United States of America 102, 6801 (2005).

21. Daniel S. Weinstock, Chitra Narayanan, Anthony K. Felts, Michael Andrec,. Ronald M. Levy, Kuen-Phon Wu, and J. Baum*, (2007), 10.1021/JA0677517.

22. R. Zhou, B. J. Berne, and R. Germain, Proceedings of the National Academy of Sciences 98, 14931 (2001).

23. G. Saladino, S. Pieraccini, S. Rendine, T. Recca, P. Francescato, F. Speranza, and M. Sironi, Journal of the American Chemical Society 133, 2897 (2011).

24. G. Bussi, F. L. Gervasio, A. Laio, and M. Parrinello, Journal of the American Chemical Society 128, 13435 (2006).

25. A. Berteotti, A. Barducci, and M. Parrinello, Journal of the American Chemical Society 133, 17200 (2011).

26. A. Ardevol, G. A. Tribello, M. Ceriotti, and M. Parrinello, Journal of Chemical Theory and Computation 11, 1086 (2015).

27. Y. Duan, C. Wu, S. Chowdhury, M. C. Lee, G. Xiong, W. Zhang, R. Yang, P. Cieplak, R. Luo, T. Lee, J. Caldwell, J. Wang, and P. Kollman, Journal of Computational Chemistry 24, 1999 (2003).

28. R. B. Best and G. Hummer, The Journal of Physical Chemistry B 113, 9004 (2009).

29. W. L. Jorgensen, J. Chandrasekhar, J. D. Madura, R. W. Impey, and M. L. Klein, The Journal of Chemical Physics 79, 926 (1983).

30. M. J. Abraham, T. Murtola, R. Schulz, S. Pall, J. C. Smith, B. Hess, and E. Lindahl, SoftwareX 1-2, 19 (2015).

31. T. Darden, D. York, and L. G. Pederson, J. Chem. Phys. 98, 952 (1993).

32. B. Hess, H. Bekker, H. J. C. Berendsen, and J. G. E. M. Fraaije, J.Comput.Chem. 18, 1463 (1997).

33. S. Miyamoto and P. A. Kollman, Journal of Computational Chemistry 13, 952, https://onlinelibrary.wiley.com/doi/pdf/10.1002/jcc.540130805.

34. H.J. C. Berendsen, J. P. M. Postma, W. F. van Gunsteren, A. DiNola, and J. R. Haak, The Journal of Chemical Physics 81, 3684 (1984).

35. G. Bussi, D. Donadio, and M. Parrinello, The Journal of Chemical Physics 126, 014101 (2007).

36. M. Parrinello and A. Rahman, Journal of Applied Physics 52, 7182 (1981).

37. J. D. Chodera, W. C. Swope, J. W. Pitera, C. Seok, and K. A. Dill, Journal of Chemical Theory and Computation 3, 26 (2007).

38. Y. Shen and A. Bax, Journal of Biomolecular NMR 48, 13 (2010).

39. D. S. Weinstock, C. Narayanan, A. K. Felts, M. Andrec, R. M. Levy, K.-P. Wu, and J. Baum, Journal of the American Chemical Society 129, 4858 (2007), pMID: 17402734, https://doi.org/10.1021/ja0677517.

40. S. Lloyd, IEEE Trans. Inf. Theor. 28, 129 (2006).

41. A. Hyv¨arinen, J. Karhunen, and E. Oja, Independent Component Analysis, Adaptive and Learning Systems for Signal Processing, Communications, and Control (John Wiley & Sons, Inc., New York, USA, 2001).

42. M. P. Harrigan, M. M. Sultan, C. X. Hernandez, B. E. Husic, P. Eastman, C. R. Schwantes, K. A. Beauchamp, R. T. McGibbon, and V. S. Pande, Biophysical Journal 112, 10 (2017).

43. M. Bonomi, D. Branduardi, G. Bussi, C. Camilloni, D. Provasi, P. Raiteri, D. Donadio, F. Marinelli, F. Pietrucci, R. A. Broglia, and M. Parrinello, Computer Physics Communications 180, 1961 (2009).

44. D. De Sancho, J. Mittal, and R. B. Best, Journal of Chemical Theory and Computation 9, 1743 (2013).

45. R. B. Best and J. Mittal, Proceedings of the National Academy of Sciences 108, 11087 (2011), http://www.pnas.org/content/108/27/11087.full.pdf.

46. A. M. Razavi and V. A. Voelz, Journal of Chemical Theory and Computation 11, 2801 (2015).

47. H. Wan, G. Zhou, and V. A. Voelz, Journal of Chemical Theory and Computation 12, 5768 (2016).

48. P. G. Bolhuis, Biophysical Journal 88, 50 (2005).

49. M. Andrec, A. K. Felts, E. Gallicchio, and R. M. Levy, Proceedings of the National Academy of Sciences 102, 6801 (2005), http://www.pnas.org/content/102/19/6801.full.pdf.

50. D. De Sancho, J. Mittal, and R. B. Best, Journal of Chemical Theory and Computation 9, 1743 (2013).

51. P. Deuflhard and M. Weber, Linear Algebra and its Applications 398, 161 (2005), special Issue on Matrices and Mathematical Biology.

52. S. R¨oblitz and M. Weber, Advances in Data Analysis and Classification 7, 147 (2013).

53. W. E. and E. Vanden-Eijnden, Journal of Statistical Physics 123, 503 (2006).

54. P. Metzner, C. Schutte, and E. Vanden-Eijnden, Multiscale Modeling & Simulation 7, 1192 (2009).

55. F. No´e, C. Schu¨tte, E. Vanden-Eijnden, L. Reich, and T. R. Weikl, Proceedings of the National Academy of Sciences 106, 19011 (2009), http://www.pnas.org/content/106/45/19011.full.pdf.

56. M. K. Scherer, B. Trendelkamp-Schroer, F. Paul, G. Perez-Hernandez, M. Ho↬mann, N. Plattner, C. Wehmeyer, J.-H. Prinz, and F. Noe, Journal of Chemical Theory and Computation 11, 5525 (2015).

57. J. Wang and A. L. Ferguson, Phys. Rev. E 93, 032412 (2016).

58. E. Schneider, L. Dai, R. Q. Topper, C. Drechsel-Grau, and M. E. Tuckerman, Phys. Rev. Lett. 119, 150601 (2017).

59. A. Mardt, L. Pasquali, H. Wu, and F. No´e, Nature Communications 9, 5 (2018).

60. J. M. L. Ribeiro, P. Bravo, Y. Wang, and P. Tiwary, The Journal of Chemical Physics 149, 072301 (2018).

